# Aryl hydrocarbon receptor and TGF-β inducible early gene mediate a transcriptional axis modulating immune homeostasis in mosquitoes

**DOI:** 10.1101/2020.11.05.368985

**Authors:** Aditi Kulkarni, Ashmita Pandey, Patrick Trainor, Samantha Carlisle, Wanqin Yu, Phanidhar Kukutla, Jiannong Xu

## Abstract

Immune homeostasis ensures effective pathogen defense and avoids overactivity, which is achieved through an orchestrated transcriptional network. Here we demonstrate that mosquito AhR and TIEG mediate a transcriptional axis to modulate the immune response. The AhR agonist compromised the immunity with reduced survival upon the challenge with bacterium *Serratia fonticola*, while the AhR antagonists enhanced the immunity with increased survival. The phenotype of pharmacological immune enhancement was corroborated genetically by the *AhR* gene silencing. The transcriptome comparison following AhR manipulations highlighted a set of AhR regulated genes, from which transcription factor TIEG, the ortholog of Krüppel-like factor 10, was chosen for further study. TIEG was required for the AhR mediated immune modulation. Silencing *TIEG* increased survival and reversed the immune suppression mediated by agonist-activated AhR. Among the transcriptomes, there were genes sharing co-expression patterns in the cohorts with AhR manipulation pharmacologically or genetically. Moreover, the mosquitoes with silenced *TIEG* and *AhR* shared ~68% altered genes upon infection. Together, the data suggest TIEG is downstream of AhR, acting as a major transcription factor mediating the immune modulation. The TIEG targets include genes involved in sugar metabolism and circadian rhythms, both processes are critical for immune homeostasis. In the naïve mosquitoes, the AhR-TIEG axis prevents the adverse effect of the overactivated IMD pathway created by silencing the inhibitor *Caspar*. In summary, AhR and TIEG constitute a transcriptional axis that mediates a gene network critical for maintaining immune homeostasis.

**Significance:** Immune homeostasis is sustained by various parameters involving different transcriptional regulatory networks. Such knowledge in mosquitoes remains scarce. Here, using AhR manipulation and transcriptome interrogation, we demonstrate that AhR and TIEG (a KLF10 ortholog) constitute a transcriptional axis to mediate immune modulation using an antibacterial immune model in the malaria vector *Anopheles gambiae*. AhR is a ligand-activated transcription factor that senses environmental signals and transcribes relevant genes to modulate immune responses. TIEG/KLF10, conserved from invertebrates to mammals, mediates various transcriptional networks. Our data show that the AhR-TIEG axis controls the genes involving in sugar sensing and circadian rhythms in the infection context. This finding warrants further study to elucidate the transcriptional control of metabolic and circadian behind immune homeostasis.

## Introduction

Mosquitoes, like many insects, have evolved an efficient innate immune system consisting of Toll, IMD, JAK-STAT, and RNAi pathways to defend against infection with bacteria, fungi, viruses, and parasites (1–4). During evolution, various regulatory circuits have evolved in insects to execute an effective defense and avoid the deleterious effects of overaggressive immune responses and immunopathogenesis(5). It has been well documented that intrinsic negative regulators Cactus, Caspar, and SOCS for the Toll, IMD, and Jak-STAT pathways, respectively, delicately tether the immune pathways. Besides, chemical defense mechanisms play important roles in homeostasis. Xenobiotic sensors are located at the interface to detect the chemicals that originated from the associated microbiota or environment. These sensors recognize ligands and initiate responses to coordinate context-dependent xenobiotic metabolism and immune defenses (6). The aryl hydrocarbon receptor (AhR) is a ligand-activated transcription factor, which was identified in the 1970s as a xenobiotic sensor recognizing the compound 2,3,7,8-tetrachlorodibenzo-p-dioxin (TCDD) in mice. Based on the studies over the past five decades, it is well known that AhR can recognize various endogenous and exogenous ligands (7) and mediate transcriptions of target genes participating in multiple physiological processes, including immune regulation and xenobiotic metabolism (8, 9). AhR is an ancient protein with an ortholog identified in the placozoan *Trichoplax* and conserved up to humans (9, 10). According to the studies in the fruit fly *Drosophila melanogaster*, the nematode *Caenorhabditis elegans*, and the cnidarian *Nematostella vectensis*, the ancestral functions of AhR are related to the development of sensory structures as well as neural systems). These ancestral functions are likely mediated by endogenous ligands. During evolution, as an add-on function, AhR has further acquired the capability to recognize ligands derived from exogenous chemicals (9, 10). Vertebrate AhR is a critical player in coordinating transcriptional circuits for immune regulation upon recognition of certain xenobiotics (11). The connections between a xenobiotic sensor and immune regulation unveil the role of chemical sensing systems in transducing non-self signals into proper responses to execute effective immune defense while maintaining homeostasis. In contrast to the rich studies about the diverse functions of AhR in the vertebrates, very little is known about the AhR in insects except for its role in development. Recently, Sonowal *et al* showed that AhR in fruit fly and nematodes can recognize indoles derived from symbiotic bacteria and initiate transcription of a gene set contributing to healthspan in the flies and nematodes (12). In this study, we demonstrate that AhR and Krüppel-like factor TIEG are connected to a transcriptional axis mediating immune modulation, suggesting that at the insect level AhR has acquired the function to modulate innate immune responses.

## Results

### Manipulation of AhR activity affects immunity against bacterial infection in mosquitoes

AhR is conserved from invertebrates to mammals. AhRs from mosquitoes *An. gambiae* and *Aedes aegypti* were clustered with the AhRs from the fruit fly *Drosophila melanogaster* and nematode *Caenorhabditis elegans* in the same clade, while AhRs from zebrafish *Danio rerio*, mouse *Mus musculus*, and human *Homo sapiens* were grouped together (Fig. S1A). The bHLH and PAS A domains are involved in DNA binding and dimerization with cofactors, which are more conserved compared to PAS B domain that is involved in ligand binding (Fig. S1B). The higher level of divergence in PAS B suggests diversity in ligand recognition. *AhR* transcripts were detected in the whole body of larvae, pupae, and adults (Fig. S1 C), indicating that *AhR* is constitutively expressed in all life forms of the mosquito. To investigate the immune regulatory role of AhR, we pharmacologically manipulated AhR activity and then examined the survival of mosquitoes upon bacterial infection. Kynurenine (Kyn) is an endogenous AhR ligand (13), which is an intermediate metabolite of tryptophan oxidation catalyzed by tryptophan-2,3-dioxygenase (TDO) in mosquitoes (14). TDO can be inhibited by 680C91(15), which leads to the reduction of endogenous Kyn production. CH233191 (16) and SR1 (15) are known AhR antagonists. After pharmacological AhR manipulation through feeding respective chemicals in sugar diet, the mosquitoes were challenged with bacteria by injecting *Serratia* into the hemocoel, and the survival rate was recorded at 24h post-injection. As shown in Fig. 1A, the vehicle control with a basal AhR activity showed a 58.6% survival. The AhR agonist Kyn reduced the survival to 39.9%. In contrast, the AhR antagonists CH233191, SR1, or 680C91 increased the survival to 78.7-85.4%. This phenotypic pattern was corroborated by the genetic manipulation of AhR. The RNAi-mediated *AhR* knockdown increased survival to 82.4% from the 57.8% in the dsGFP control (Fig.1B).

**Figure 1.**
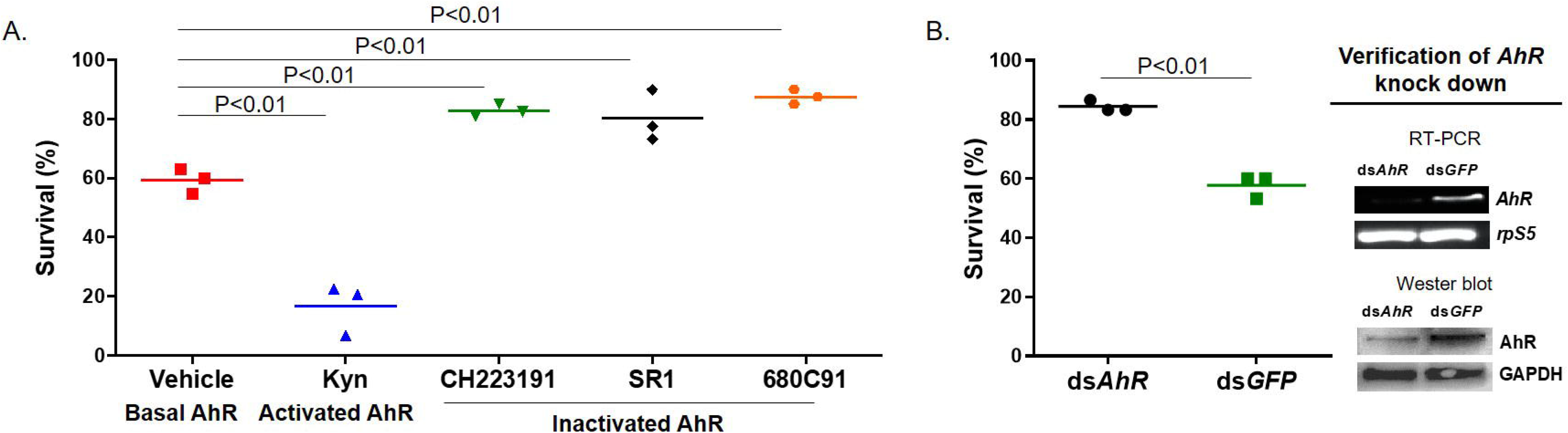
Mosquito survival upon bacterial challenge when AhR activity is manipulated pharmacologically or genetically. **(A)** The effect of pharmacological manipulation of AhR activity on the survival post *Serratia* challenge. Average survival (%) was denoted by the line. **(B)** The effect of *AhR* silencing on the survival post-challenge. The *AhR* knockdown was verified by RT-PCR and Western blot. Survival data were generated from triplicate experiments and compared by Chi-square.

### Transcriptomic alterations upon AhR manipulation

To identify genes that are regulated by AhR upon bacterial infection, we conducted RNA sequencing (RNA-seq) to profile the transcriptomes upon the treatment. Mosquitoes were fed with AhR antagonist SR1 or vehicle control and then challenged with *Serratia*. Sterile water injection was used as injury control. Surviving mosquitoes at 24hr post-challenge were processed for RNA-seq. The transcriptional abundance of genes was measured using normalized read counts, and differentially expressed genes were identified by DESeq2. Between the *Serratia* infection and injury control, 2102 genes were differentially expressed (using a cutoff of q-values <0.05, Table S3), which we defined as infection responsive genes. Among these genes, 265 were upregulated and 145 were downregulated at least 2-fold. The infection inducible gene set includes typical immune genes, such as immune pathway components (spaetzle1, spaetzle3, PGRPLB, Rel1, and Rel2), immune effectors (DEFs, CECs, TEPs and LRIMs, CLIP serine proteases, fibrinogens), and immune regulators, such as serpins and inhibitors of apoptosis (IAPs). The AhR antagonist affected the expression of 696 infection-responsive genes, among them 66 upregulated and 36 downregulated genes had at least 2-fold differences. The AhR antagonist affected genes are largely unannotated, and the major immune genes are not affected. The expression patterns of *Def1, Tep15, SRPN10*, and *TIEG* were validated by qRT-PCR (Fig. S2).

### AhR and TIEG mediate a transcriptional axis in the infection context

One of the AhR regulated genes, *AGAP009889*, is the ortholog of *cabut* in Drosophila, which was named as TGF-β-inducible early-response gene (TIEG) (17) after the human homolog (18). Drosophila TIEG is involved in the development, metabolism, and growth control (19–22). TIEG is a zinc finger transcription regulator in the Sp1-like/Krüppel-like family, designated as Krüppel-like factor (KLF) 10 (23). TIEG/KLF10 is involved in the TGF-β/Smad signaling (24–26). AhR and TGF-β/Smad signaling are connected to mediate immune modulation to maintain immune homeostasis in mice (27, 28). Therefore, we tested the hypothesis that TIEG was downstream of AhR to regulate the transcription of genes responsible for immune modulation. First, the putative AhR binding motif (GCGTG) (29) was identified in the sequence upstream of the TIEG coding region, suggesting *TIEG* may be an AhR target gene. Next, we examined the effect of dsTIEG on the *Serratia* infection outcome. As expected, dsTIEG resulted in 90.1% survival, higher than 55.7% in the dsGFP control (Chi-square test, P<0.01, Fig. 2A), a pattern similar to the dsAhR effect. Then, we conducted RNA-seq to compare the transcriptomes in response to the dsAhR and dsTIEG and AhR antagonist. Clustering analysis separated transcriptomes into two clusters (Fig. 2B). The injury control was separated from the *Serratia* infection and the antagonist/infection in one cluster. In the other cluster, the dsAhR/infection and the dsTIEG/infection were grouped and distinct from the dsGFP/infection. The PCA analysis revealed that principal component 1 (PC1) and PC2 explained 61% and 11.2% of the variance, respectively, the transcriptome replicates of each treatment were closely clustered, and the transcriptomes of different treatments were separate distinctly (Fig. 3). Replicates of dsAhR and dsTIEG were located closely, indicating a similarity in their transcriptomic response. However, the replicates of dsGFP/infection were separate distinctly from the replicates of *Serratia* infection, suggesting that the dsGFP treatment had other effects. In addition, there was an evident separation between the AhR antagonist and the *AhR* silencing, suggesting that each of the two approaches had other effects in addition to affecting AhR activity. The AhR antagonist was administrated via diet, while the dsRNA was administered through injections, an invasive method. In insects, dsRNA triggers defense against viruses (30, 31). The dsGFP injection may have an unknown influence on the response to the following bacterial challenge. Indeed, in response to the *Serratia* challenge, the cohorts with naïve backgrounds and the cohort with dsGFP treatment shared only 450 genes, and the dsGFP cohorts demonstrated 2099 unique responsive genes, and the native cohorts had 481 unique responsive genes (Fig. S3). These confounding effects may contribute to the patterns revealed by PC1 and PC2. The PC3 and PC4 captured 7.3% and 4.1% of the variance, the replicates of AhR-antagonist/infection, dsAhR/infection and dsTIEG/infection were clustered more closely, which separated from the clusters of *Serratia* infection and dsGFP/infection (Fig. 3). Therefore, PC3 and PC4 may represent the effect of AhR manipulation more genuinely and describe more accurately the transcriptomic responses that are driven by the AhR-TIEG signaling axis. This pattern was further corroborated by co-expression module analysis. The weighted gene correlation network analysis (WGCNA) was implemented to identify phenotype-associated modules of co-expressed genes. WGCNA identified 35 modules from the entire dataset, each containing 23-1382 differentially expressed (DE) genes. The treatment-affected genes were not distributed evenly in these modules. As shown in Fig. S4. Category I includes 9 modules, representing 62.1% (1305/2102) DE genes in the infection cohorts, but only 30.5% (498/1631) in the AhR antagonist cohorts, 30.6% (1007/3285) in the dsAhR cohorts, and 27.2% (888/3260) in the dsTIEG cohorts. Category II includes 5 modules, representing 8.1% (170/2102) DE genes in the infection cohorts, but 26.3% (429/1631) in the antagonist, 37.2% (1223/3285) in the dsAhR and 36.4% (1185/3260) in the dsTIEG cohorts. Overall, the modules in Category I and II represent genes targeted by the AhR-TIEG axis (Fig. S4). Together, the transcriptome comparisons indicate that the two manipulation approaches had resulted in broad impacts on a variety of genes with different functions. The PCA and WGCNA module analysis effectively extracted transcriptomic patterns caused by the AhR manipulations. The transcriptomic pattern was validated by qRT-PCR, the transcription of *SRPN10, TEP15, SOCS* was down-regulated by *AhR* and *TIEG* silencing in response to the bacterial challenge (Fig. 2C). Next, we reasoned that the immune suppression resulted from AhR activation would be attenuated by *TIEG* silencing. Indeed, the survival reduction by Kyn feeding was reversed by dsTIEG (Chisquare, P<0.01, Fig. 2D). This result suggests that TIEG acts downstream of AhR. This conclusion was corroborated by the transcriptional patterns in the dsAhR and dsTIEG cohorts. As shown in Fig. 4A, 3285 genes and 3256 genes in the dsAhR and the dsTIEG cohorts were altered upon the *Serratia* challenge, respectively. Among them, 2245 genes were shared by both treatments, accounting for 68.3% and 68.9% of the total genes that were affected in the context, and these genes exhibited the same transcriptional directions in the dsAhR and dsTIEG cohorts (Fig. 4B). This remarkable pattern indicates that approximately 2/3 of the AhR-regulated genes are TIEG target genes, supporting the hypothesis that AhR and TIEG act as a transcriptional axis in response to the infection. TIEG/KLF10 can be a transcriptional activator or repressor (32). This pattern remains in the mosquitoes as well. In the dsAhR and dsTIEG cohorts, the expression was enhanced in 1150 genes and was decreased in 1095 genes, respectively, indicating that the AhR-TIEG axis represses or enhances the transcription of these genes. The mammalian KLF10 and Drosophila TIEG (Cabut) are involved in the ChREBP/Mondo-Mlx transcriptional network for sugar sensing, which is connected to the circadian rhythm (20, 33). In this context, KLF10/TIEG is a transcriptional repressor of phosphoenolpyruvate carboxykinase (PEPCK). In the mosquitoes, *pepck* was induced by the infection, but this induction was under the repressive control of the AhR-TIEG axis as the *pepck* expression was increased in both dsAhR and dsTIEG upon the *Serratia* challenge (Fig. S5). In Drosophila, sugarbabe is downstream of Mondo-Mlx and responsible for lipid homeostasis (34). *AGAP006736*, the ortholog of *sugarbabe*, was increased in both dsAhR and dsTIEG cohorts, suggesting that its expression was repressed by AhR-TIEG (Fig. S5). The circadian clock orchestrates immunity (35) as well as metabolism (36). In the dsAhR and dsTIEG cohorts, circadian genes *tim, cwo*, and *vri* were upregulated, suggesting they were negatively controlled by the axis (Fig. S5). A set of immune genes were repressed by the axis. For instance, *Toll5A, Rel2, ClipB19, TEP3* and *TEP15.* These genes were all induced by the infection, but their transcription level was capped under the control of the axis. The axis acts as a transactional activator as well. Silencing the axis reduced the transcription of a set of genes, including Mondo-Mlx target genes *arrestin 1* and *2, G-protein coupled receptors* (*GPRop1* and *3*), and *vitellogenin receptor* (Fig. S5).

**Figure 2.**
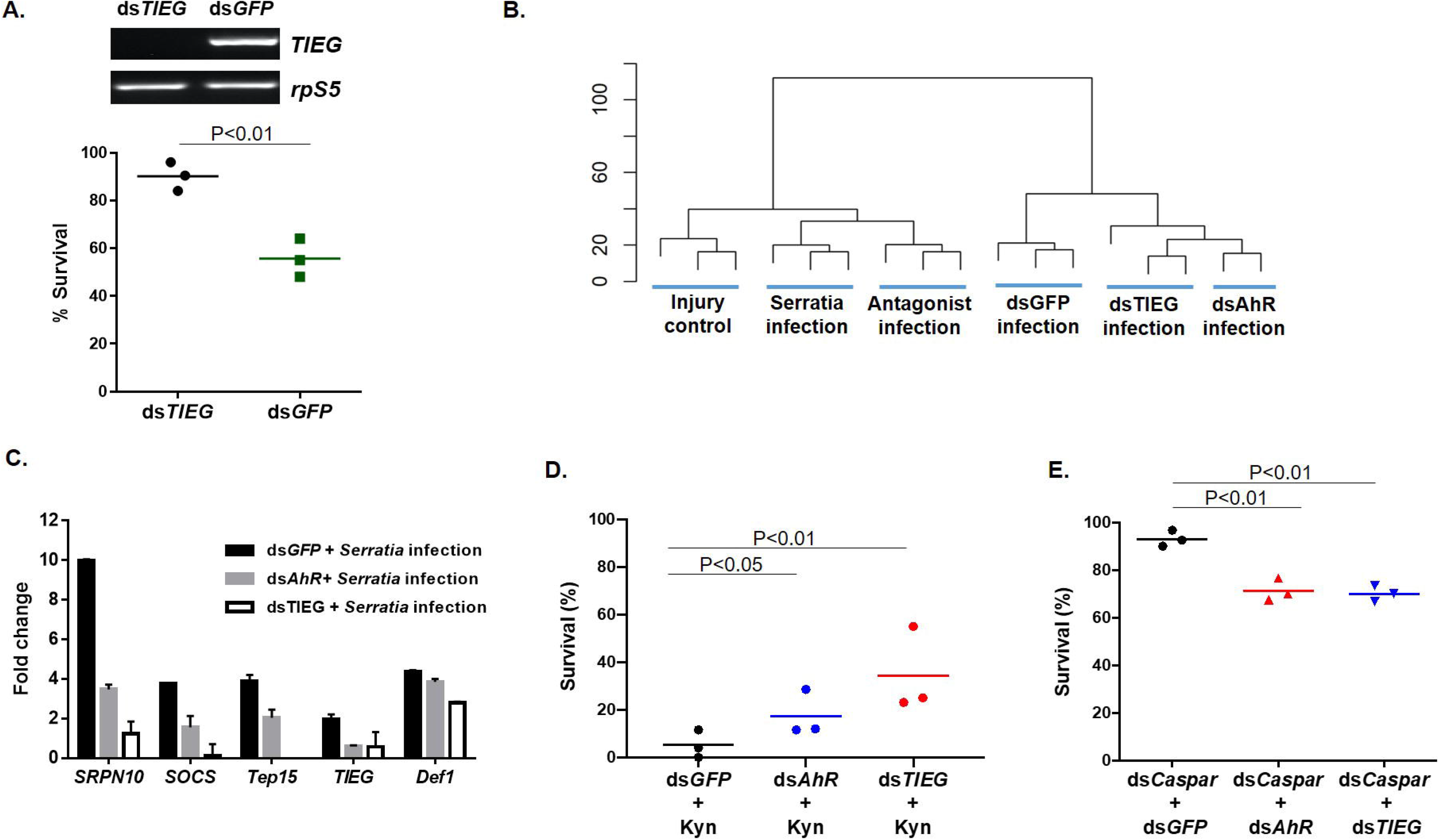
TIEG is mediating immunomodulation downstream of AhR. **(A)** dsTIEG increased the survival upon *Serratia* challenge. The *TIEG* knockdown was verified by RT-PCR. **(B)** Clustering of transcriptomes in response to *Serratia* challenge. The scale bar represents the distance between the clusters. **(C)** qPCR validation of selected target genes of AhR-TIEG axis. **(D)** The Kyn-mediated immune suppression was reversed by dsAhR and dsTIEG. **(E)** The effect of dsAhR and dsTIEG on the survival of mosquitoes in which the IMD was overactivated by dsCaspar. Survival data were generated from triplicates and compared by the Chi-square test.

**Figure 3.**
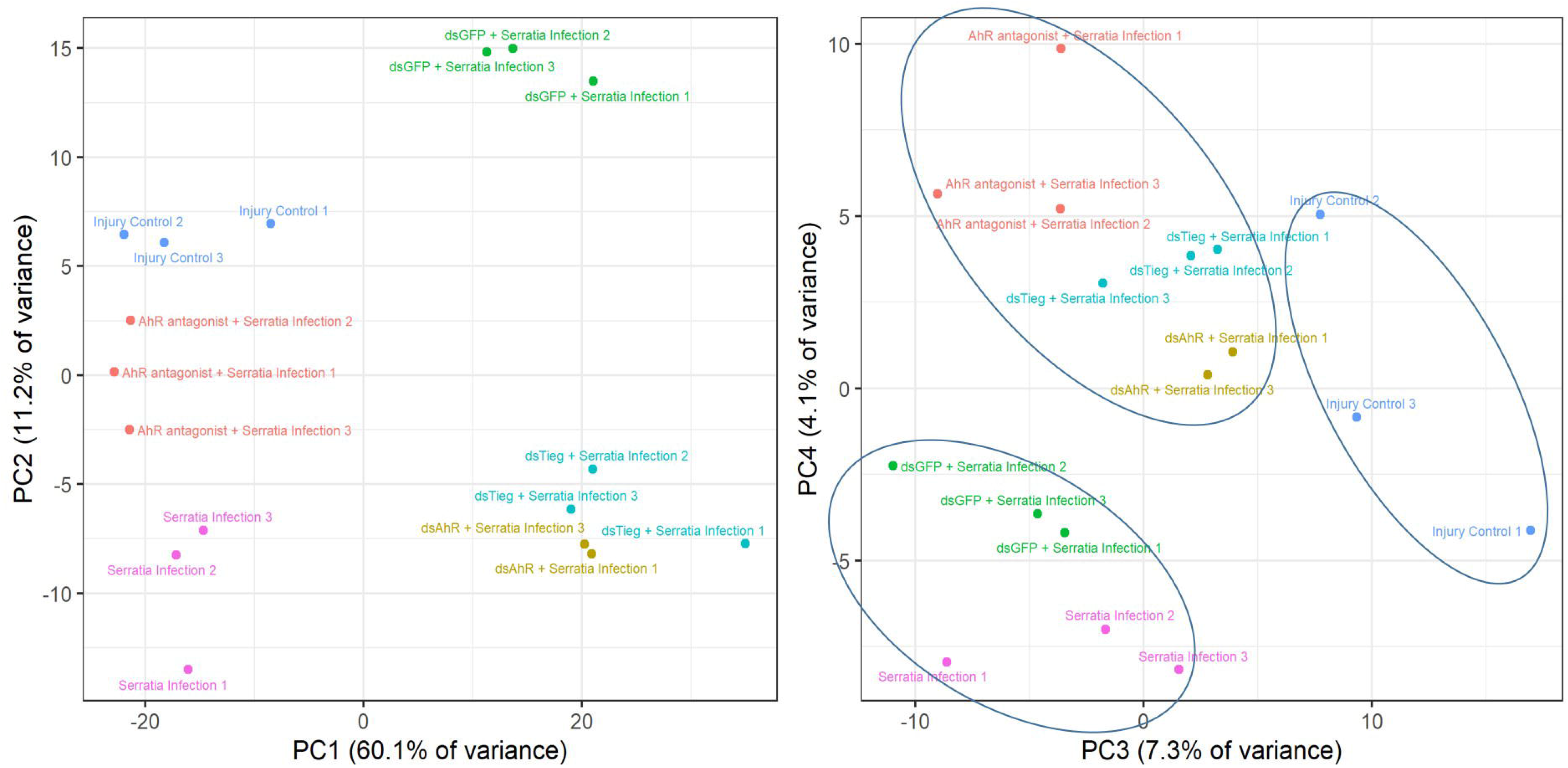
Principal component analysis (PCA) of transcriptomes. The PC1 and PC2 explained the major variance of transcriptomes with different treatments. The impact of AhR antagonist, dsAhR and dsTIEG on the transcriptomic response to the challenge was better represented by PC3 and PC4.

**Figure 4.**
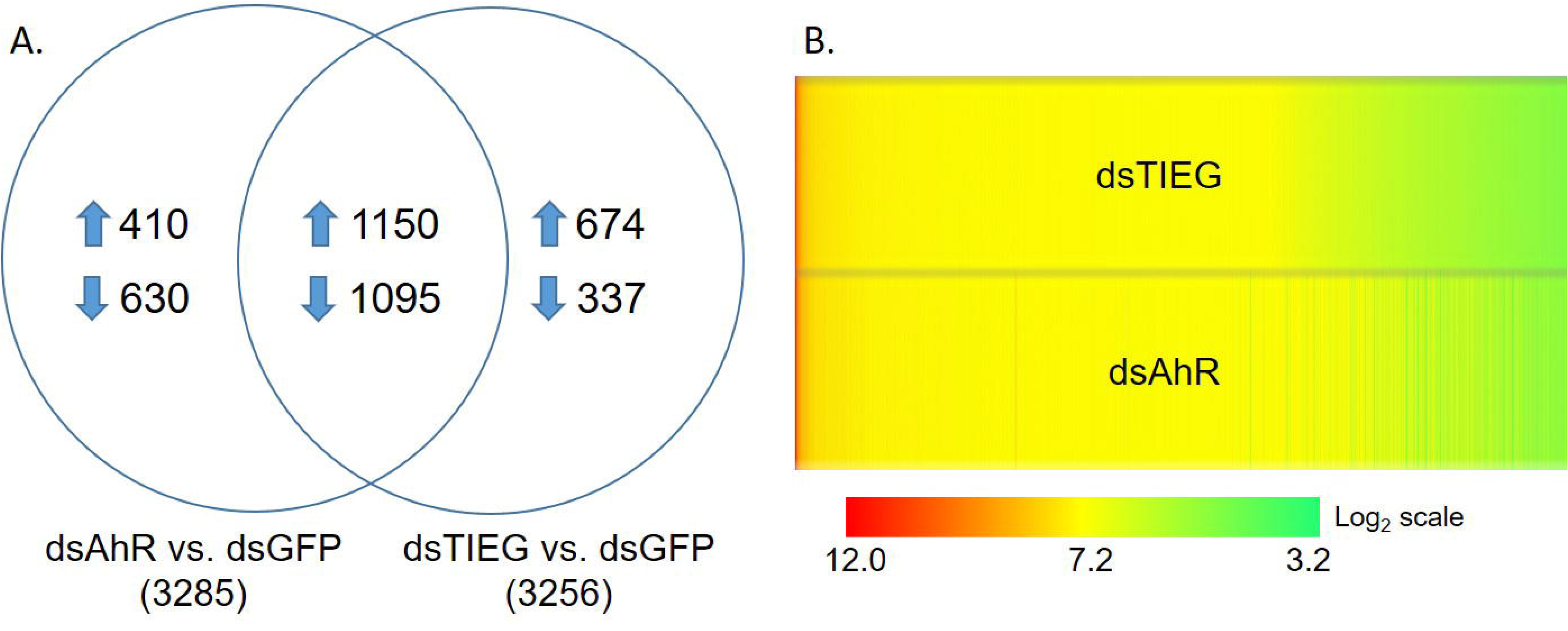
Expression patterns of the genes that were regulated by the AhR-TIEG axis. **(A)**The distribution of altered genes by respective treatments. **(B)** The heatmap of transcript abundance (TPM) of the genes shared in the dsAhR and dsTIEG cohorts.

### The AhR-TIEG axis counteracts immune overactivation

From the observations above, we conclude that AhR and TIEG constitute a signaling axis to modulate the immune response. This axis may help maintain immune homeostasis by counteracting adverse effects of immune overactivation. To test this hypothesis, we created an over-activated immune state by silencing *Caspar*, the negative regulator of IMD pathway (37), and examined the impact of *AhR* or *TIEG* silencing on the survival curve. As shown in Fig. 2E, the cohorts of dsGFP, dsAhR/dsGFP, dsTIEG/dsGFP, and dsCaspar/dsGFP remained ~90% survival at day 3 post respective injections. However, when *Caspar* was co-silenced with either *AhR* or *TIEG*, the survival was reduced to 70%. The results show that the overactivation of IMD by depleting suppressor *Caspar* in the absence of an immune insult displays a normal survival only when AhR and TIEG are present, suggesting that the adverse effect of IMD overactivation is prevented by the AhR-TIEG axis.

## Discussion

AhR is conserved from invertebrate to vertebrate. During evolution, in addition to its ancestral role in the development of sensory structure and neural systems, the AhR has diversified into a chemical sensor with binding affinity for a broad spectrum of intrinsic and/or extrinsic ligands derived from the environment or associated microbiota (38). The ligand recognition transduces the chemical signals into responses in various contexts, such as xenobiotic detoxification and immune regulation in vertebrates. In *C. elegans* and *D. melanogaster*, triggered by bacterial indoles, AhR directs a process to extend healthspan (12). In this study, we demonstrate that mosquito AhR and TIEG constitute a transcriptional axis that directs immune regulation. We examined the role of AhR in antibacterial immunity. The AhR activation by agonists reduced the immunity, while the AhR inactivation by antagonists or the TDO inhibitor enhanced the immunity (Fig. 1A). In line with the pharmacological evidence, silencing *AhR* increased immunity as well (Fig. 1B). The AhR target genes in the context were screened by transcriptome interrogation. A variety of genes were affected by the AhR manipulations, including several transcription factors. We further characterized a C2H2 transcription factor, TIEG, which is the vertebrate ortholog of Krüppel like factor 10 (39). Drosophila and vertebrate TIEG is involved in the TGF-β1 pathway (22, 26). TGF-β signaling plays a critical role in immune regulations in both invertebrates and vertebrates (40). In *Drosophila*, upon injury and bacterial infections, two TGF-β signals in hemocytes are induced by respective cues, BMP (*dpp*) represses antimicrobial peptide production, and Activin (*daw*) suppresses melanization (41). Recently, AhR-TGF-β1/Smad signaling has been shown to mediate immune suppression in scurfy mice, a mouse autoimmune model (27). We generated three lines of evidence to demonstrate that the TIEG is downstream of AhR to mediate the immune regulation. First, *TIEG* silencing enhanced antibacterial immunity (Fig. 2A); second, the immune suppressive effect induced by the AhR activation via Kyn was reversed by dsAhR and dsTIEG, respectively (Fig. 2D); third, dsAhR and dsTIEG resulted in similar transcriptional profiles, dsAhR affected 2245 (~68 %) genes were altered by dsTIEG as well (Fig. 4). The AhR-TIEG axis appears to function as a negative regulatory loop that sustains a homeostatic immune state. As shown in Fig. 2E, the AhR-TIEG axis is required for suppressing the deleterious effect of dsCaspar in naïve mosquitoes. Taken together, AhR and TIEG mediate a transcriptional network to carry out immunoregulatory functions.

It is long known that vertebrate AhR mediates immunomodulation through its target genes (11). However, many AhR target genes can also be regulated by other transcription factors, it is challenging to characterize these genes in the immune context. In this study, TIEG was identified as a major transcription factor downstream of AhR in response to bacterial infection. The transcriptome interrogation revealed a gene network that is controlled by the AhR-TIEG axis. The TIEG/KLF10 connects energy metabolism and the circadian clock through the sugar sensing pathway (20, 33, 42). This network appears to be active in the infection as the TIEG target genes include *pepck, arrestin 1* and *arrestin 2* in the sugar sensing pathway as well as the circadian genes *tim, vri* and *cwo* (Fig. S5). Pepck converses oxaloacetate (OAA) to phosphoenolpyruvate (PEP), which is a rate-limiting step in gluconeogenesis, representing a critical link between biosynthesis and cataplerosis of the citric acid cycle (43, 44). Glycolysis and gluconeogenesis are delicately balanced as energy supply for fueling immune activities (45). Evidence is mounting that metabolic homeostasis is critical for sustaining appropriate immune functions (46, 47), and TIEG/KLF10 is a key player in metabolic regulation (48). In summary, we identified an AhR-TIEG mediated transcriptional network that orchestrates immune modulations integrating various processes, including sugar sensing and central carbohydrate metabolism. This suggests that mosquito AhR and KLF10 had acquired functions pertinent to immune homeostasis. Further studies are required to identify AhR ligands that initiate the axis and elucidate the functional connection between energy metabolism and immune response in mosquitoes.

## Materials and Methods

### Mosquitoes

*Anopheles gambiae* G3 strain was used for this study. The G3 mosquitoes were reared in an insectary with 28°C and 80% humidity, a light/dark cycling of 14:10 hours. Larvae were cultured in water pans with food (1:1 mix of the ground pellet of cat food and brewer yeast), and adults were maintained on 10% sucrose sugar meal, and fed on mice to acquire blood for egg production (animal protocol was reviewed and approved by the NMSU IACUC).

### Phylogenetic inference using AhR protein sequences

The AhR protein sequences of representative organisms from invertebrates to mammals (Table S1) were used for inferring phylogenetic relationships. The sequences were aligned using the MUSCLE algorithm. A phylogenetic tree was made by the Neighbor-Joining method using Jones-Taylor-Thornton (JTT) model, with 500 bootstrap replications. Similar tree topology was generated by the Maximum Likelihood method using the JTT model with 500 bootstrap replications. Only the NJ tree is shown in Fig. S1. The domains were visualized in Simple Module Architecture Research Toll (SMART, http://smart.embl-heidelberg.de/), and the peptide sequences of domains bHLH, PAS A, and PAS B were extracted and protein sequence identity between these organisms were compared.

### Pharmacological manipulation of AhR activity in mosquitoes

AhR antagonists and agonists were used to manipulate AhR activity in mosquitoes. In each case, chemicals were fed to newly emerged adult mosquitoes, 50-60 females per cohort, in 10% sucrose solution for three days and then challenged with *Serratia fonticola* S1 (see below). Kynurenine (Kyn) is an endogenous ligand of AhR (13). Kyn is generated during the oxidation of tryptophan, a reaction catalyzed by tryptophan-2,3-dioxygenase (TDO) in mosquitoes (14). Kyn at 30μM (Sigma-Aldrich, K8625) was used as an agonist to enhance the AhR activity. We used two AhR antagonists CH223191 at 90 μM (Sigma-Aldrich, C8124) and Stem Regenin (SR1) at 3μM (Selleckchem, S2858). Additionally, we used 680C91 (20μM) (Sigma-Aldrich, SML0287), a TDO inhibitor, to reduce endogenous Kyn production. The concentrations of these chemicals were chosen empirically based on the resulting phenotypes after administration.

### Bacterial infection

Bacterium *Serratia fonticola* S1 was isolated from a wild-caught specimen of *Aedes albopictus*, collected in Florida in July 2015. The bacteria can cause acute hemocoelic infection after intrathoracic injection (49). The bacteria were transformed with a plasmid expressing GFP as described previously (50). The bacteria were grown overnight in Luria Bertani broth containing ampicillin (100μg/ml) at 28°C. The bacterial culture at OD_600nm_=1 was diluted 1000 times with sterile H_2_O, which yielded approximately 1000 colony forming units (CFU) /μl, and approximately 100 nl of this bacterial solution was injected intrathoracically into the hemocoel of *An. gambiae* on day 3 after AhR activity was manipulated as described above. Survival at 24 hours post-infection was used to assess the antibacterial immunity as described in (49). The single thorax of dead and surviving mosquitoes was homogenized in 50 μl sterile water, and 30 μl homogenates were spread to an LB plate with Ampicillin and cultured at 28 °C overnight. The colonies on the plate were examined under UV light to visualize GFP-tagged bacteria. In the *Serratia*-injected mosquitoes, GFP-tagged bacteria were recovered, while in the mosquitoes injected with sterile water, no GFP-tagged bacteria were detected (data not shown). The data were generated from three experimental replicates. The survival between the treatment and vehicle control was tested using a Chi-square test.

### RT-PCR

Mosquito RNA was extracted from 15 females using Trizol reagent (Invitrogen, Cat# 15596026). Genomic DNA contamination was removed by DNaseI treatment using TURBO DNA-free Kit (Invitrogen, AM1906). The cDNA synthesis was carried out using NEB ProtoScript II Reverse Transcriptase (NEB, M0368S). The cDNA was used as a template for RT-PCR, to determine transcript abundance for various genes. The primers used are listed in Table S2. No reverse transcriptase (NRT) and no template control (NTC) served as negative controls.

### RNAi mediated gene knockdown in mosquitoes

RNAi-mediated *AhR, TIEG*, and *Caspar* knockdowns were performed. For dsRNA preparation, a target gene fragment was PCR amplified using gene-specific primers with the T7 promoter sequence at the 5’ end. The PCR products were used to synthesize dsRNA using a T7 RNA Polymerase Kit (Sigma-Aldrich RPOLT7-RO ROCHE). Generated dsRNAs were treated with TURBO DNA-free Kit (Invitrogen, AM1906) to remove DNA. The dsRNA of a GFP fragment was used as control dsRNA. Mosquitoes were injected with 0.5μg/μl dsRNA to initiate RNAi. To co-silence *Caspar* along with either *AhR* or *TIEG*, respective dsRNA, each at 1.0μg/μl, were mixed for injection. Newly emerged *An. gambiae* female mosquitoes, 60 females per cohort, were subjected to intrathoracic injections with ~100nl of dsRNA. Treated mosquitoes were maintained on 10% sucrose solution for three days. For the bacterial challenge, cohorts of dsGFP control, dsAhR, and dsTIEG were injected with *Serratisa* at day 4 post dsRNA treatment as described above. The gene knockdown efficacy was verified by RT-PCR with primers in Table S2.

### AhR Western blot

Rabbit Polyclonal antibody against AhR was made at GenScript (New Jersey). The antigen was a peptide fragment in the PAS B domain of AhR protein (aa 208-394). For Western blot, mosquito proteins were extracted using Insect Cell-PE LB™ (G Biosciences, 786-411) at three days post dsRNA injection. The protein samples (20 μg/well) were separated by SDS-PAGE gel and transferred to a nitrocellulose membrane. The membrane was probed by the AhR antibody (1:1000). GAPDH antibody (GeneTex, GTX100118) was used as a loading control. HRP labeled secondary antibody was used to detect signal using KPL LumiGLO kit following manufacturer’s instruction. The blot was scanned to visualize the signal using the IMAGE STUDIO, Version 5.x, LI-COR.

### Transcriptome analysis

RNAseq was used to compare transcriptomes between samples with AhR manipulation. To examine the effect of pharmacological AhR inhibition, three cohorts of females were used. Mosquitoes were fed with AhR antagonist SR1 or vehicle control for three days, and then were challenged with *Serratia* infection, as described above. An injury control (in which mosquitoes were injected with sterile H_2_O) was included to control the effect of damage associated with the injection. To examine the effect of gene knockdown, three cohorts of newly emerged females were used; each was treated with dsAhR, dsTieg, or dsGFP control, for 3 days, and then challenged with *Serratia* infection. Total RNA from 15 live mosquitoes at 24hr post-challenge was isolated using Trizol reagent, and then DNaseI treatment was followed to remove DNA contamination. Triplicate RNA samples were prepared for each treatment. The RNA samples were further processed at Genewiz, NJ, where the cDNA libraries were prepared and sequenced using Illumina Hiseq, 2 x 150 bp paired-end chemistry. At least 25M clean reads were generated from each RNA sample, which provided a sequencing depth sufficient for transcriptome analysis. The reads were mapped against transcripts of *An. gambiae* (NCBI), implemented by using Array Star v.12 (DNAstar). Read counts were normalized using the median of ratios method (51) using DESeq2 software (52). In determining normalized read counts, this method accounts for sequencing depth and RNA composition by calculating normalization factors for each sample in comparison to a pseudoreference sample. After determining normalized read counts, an independent filter was utilized which removed transcripts with normalized counts less than 5. This resulted in a dataset of 10,714 transcripts. The clustering of all samples revealed that replicate 2 of dsAhR/*Serratia* infection was not consistent with the other two replicates, likely due to a quality issue associated with the replicate, therefore, this replicated was removed from the analysis. Differentially expressed genes were identified using a negative binomial generalized linear model (GLM) available through DESeq2 (52). Likelihood ratio tests were conducted to identify transcripts that exhibited differential expression between all groups. Pairwise differential expression comparisons were made and statistical significance was determined by computing q-values that preserve the False Discovery Rate (FDR) (51, 53, 54). For example, concluding that a transcript was differentially expressed between two groups with a q-value of 0.05 would imply that there was a 5% chance (expected) that this conclusion was a false positive. To determine a lower-dimensional representation of the transcriptomic data, principal components analysis (PCA) was conducted using regularized log-transformed (rlog) data. PCA seeks to find a small set of “principal components” that capture a large proportion of the variance in the original data (55). The rlog data was determined using DESeq2, while the “prcomp” function in R (56) was utilized to determine the PCA. The proportion of the variance captured by each of the principal components was determined. Weighted gene coexpression network analysis (WGCNA) (57) was conducted to identify modules (or sets) of transcripts that are co-expressed. This analysis was conducted as follows. First, the topological overlap between transcripts in a signed and soft-thresholded correlation network determined from the rlog data was determined using the R package WGCNA (58). Further, transcript modules were determined utilizing hierarchical clustering with an “average” link function and utilizing the hybrid version of the “Dynamic Tree Cut” algorithm (59). To validate expression patterns revealed by RNAseq, a selected set of genes was measured using quantitative RT-PCR. Primers were presented in Table S2. The submission of the RNAseq reads to NCBI is in the process. TPM (Transcripts Per kilobase Million) was used for the comparison of transcriptional abundance between different conditions (60). The KEGG pathway analysis was implemented at Pathview Web, an online tool for pathway-based data integration and visualization (61). The normalized read counts were used as input for Pathview analysis. The RNA-seq datasets are available at NCBI SRA under BioProject PRJNA691571.

## Acknowledgments

We thank Dr. Jennifer Curtiss in Biology Department at New Mexico University for her valuable suggestions along the course of the study. We thank Dr. Michelle Riehle in Microbiology and Immunology at Medical College of Wisconsin for her constructive comments on the manuscript. This work was supported by the National Institutes of Health [SC1AI112786 to J.X.] and the National Science Foundation [No. 1633330 to J.X.]. The content is solely the responsibility of the authors and does not necessarily represent the official views of the National Institutes of Health.

---

**Fig. S1.**
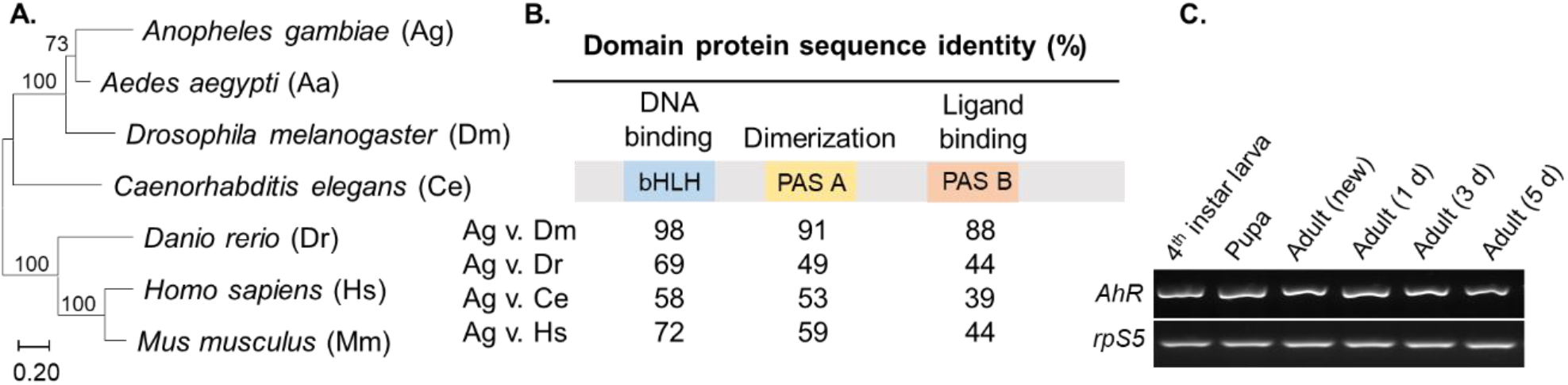
**(A)** The AhR phylogenetic tree inferred from amino acid sequence comparison using Neighbor-Joining method. Bootstrap values were displayed on nodes. The scale bar indicates the genetic distance. **(B)** The level of conservation of domains bHLH, PAS A and PAS B, represented by the identity of the protein sequence. **(C)** *AhR* is constitutively expressed in larvae, pupae and adults (newly emerged, 1-, 3-, 5-day old), assayed by RT-PCR, *rpS5* was used as cDNA input control.

**Fig. S2.**
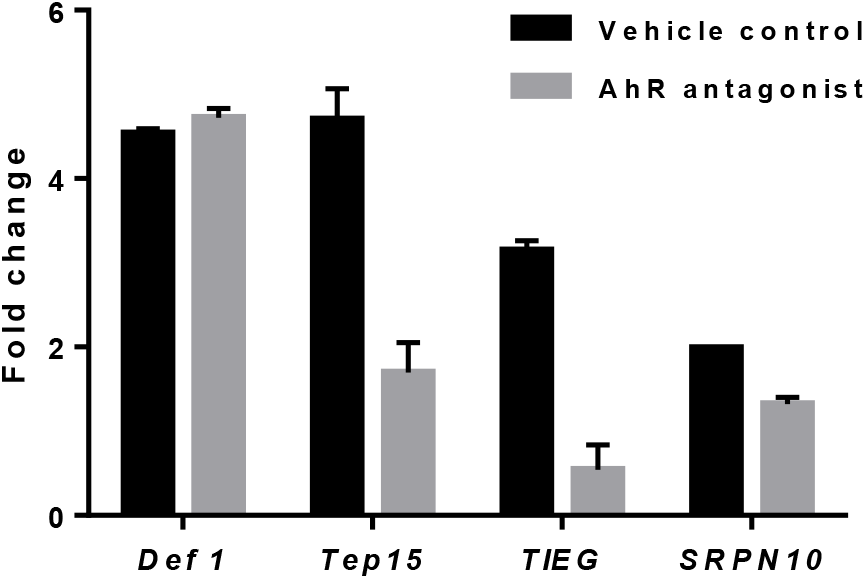
Transcriptional validation of selected genes by qRT-PCR. Def1 represents an AhR independent immune gene, and *Tep15, TIEG* and *SRPN10* represent the AhR regulated genes. Fold change was calculated relative to injury control. Error bars represent SE.

**Fig. S3.**
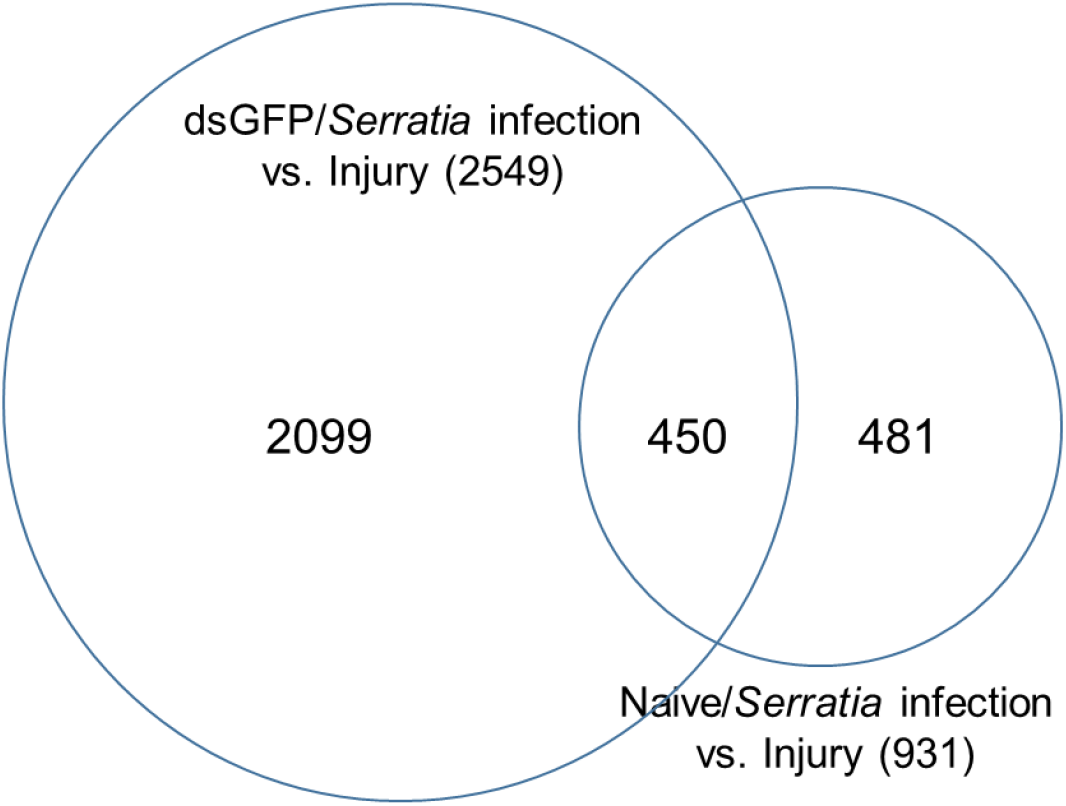
Different responses to the *Serratia* challenge between the dsGFP and naïve cohorts. The dsGFP treatment altered 2.73-fold (2549/931) more genes than the naïve background, and only 450 genes were shared between.

**Fig. S4.**
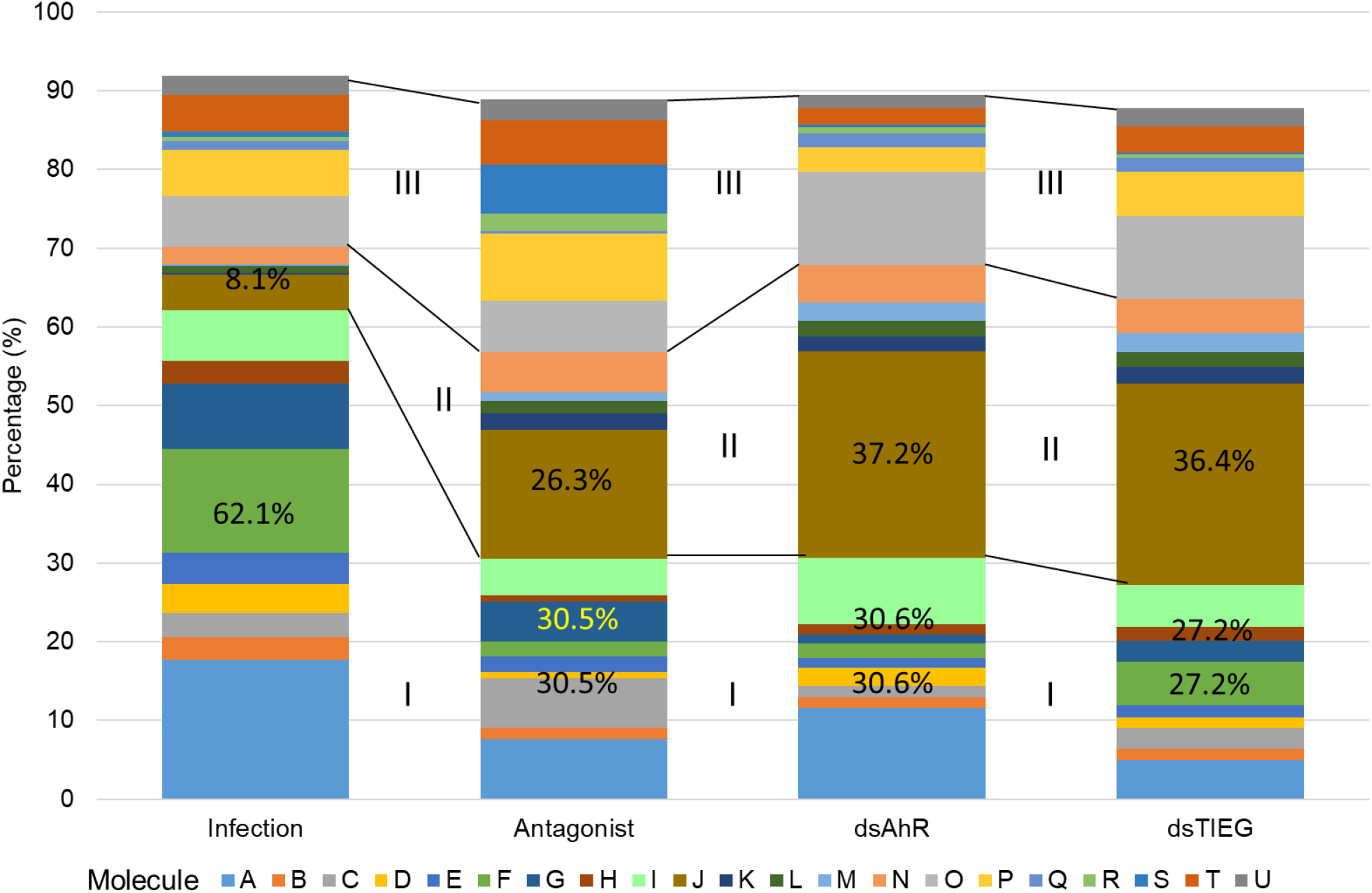
Co-expression modules analysis. The co-expression pattern was similar between the AhR antagonist, dsAhR and dsTIEG cohorts. The Category I modules contain more genes in the Infection cohorts than the AhR inactivated cohorts, while the Category II modules contain more genes in the AhR inactivated cohorts. The patterns represent the shared effects of AhR manipulation by both pharmacological and genetic approaches.

**Fig. S5.**
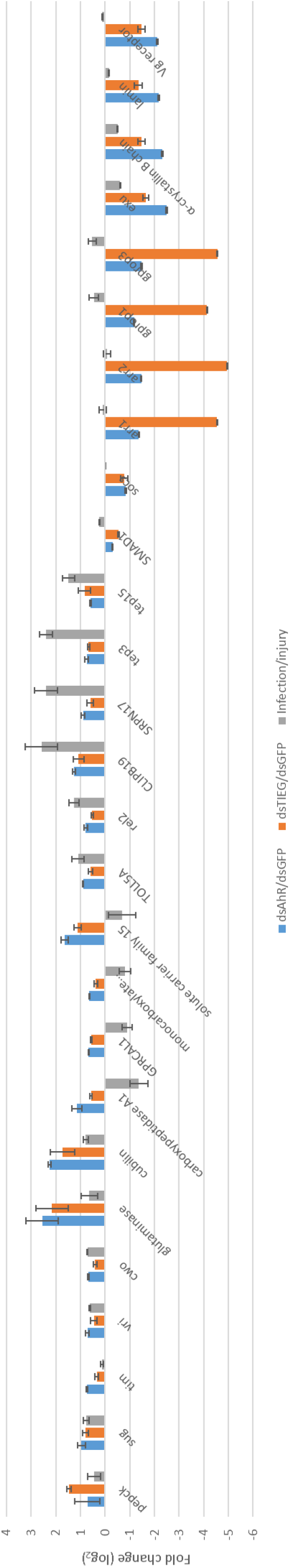
The AhR-TIEG axis-target genes. The fold change (log_2_) was calculated based on the TPM values between the dsAhR or dsTIEG and dsGFP cohorts upon the *Serratia* challenge. The fold change of infection/injury in the naïve background was presented as a reference. Bars represent standard error derived from three replicates.

## Notes

### Competing Interest Statement

The authors have declared no competing interest.

### Summary of Updates

A subset of target genes regulated by the Ahr-TIEG axis was unveiled by deeper RNA-seq data analysis. These genes are involved in sugar-sensing and circadian rhythm, which connects the transcriptional network to the metabolic and circadian regulation in immune homeostasis in mosquitoes.

