## Supplementary figures and images for "Aryl hydrocarbon receptor and TGF-β inducible early gene mediate a transcriptional axis modulating immune homeostasis in mosquitoes"

### Fig S1

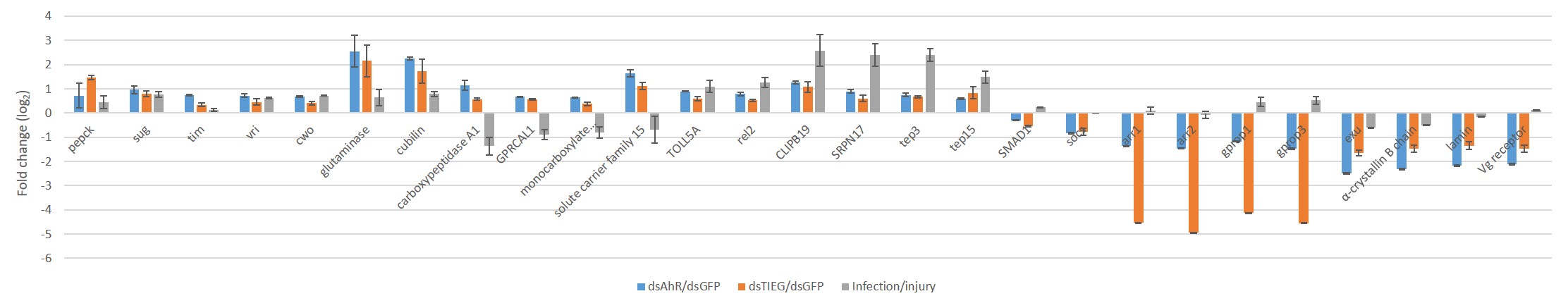

### Fig S2

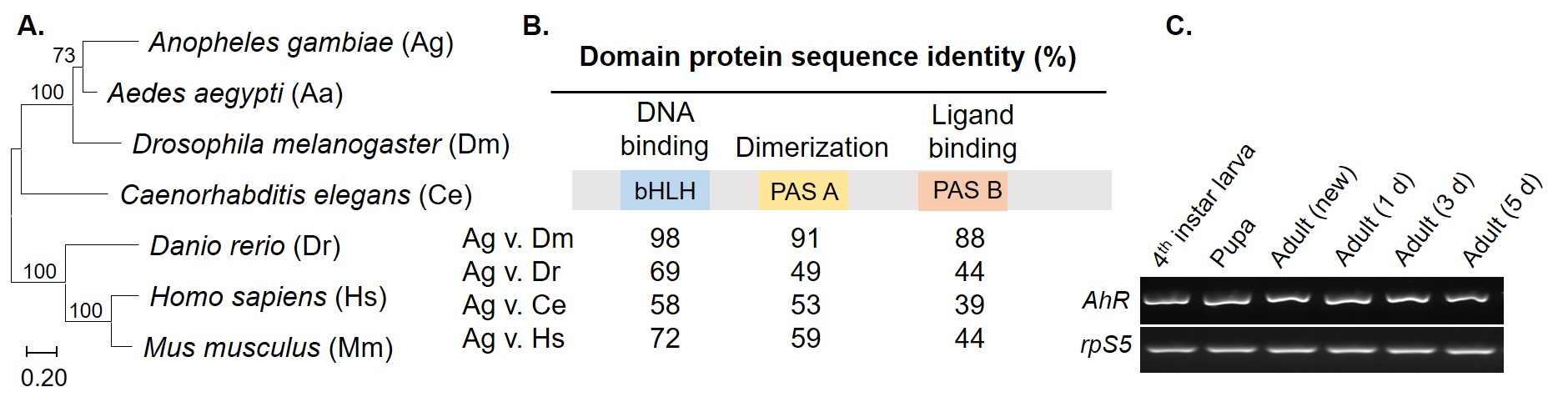

### Fig S3

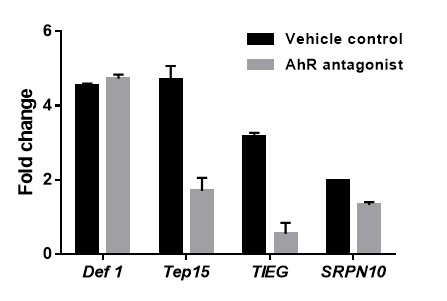

### Fig S4

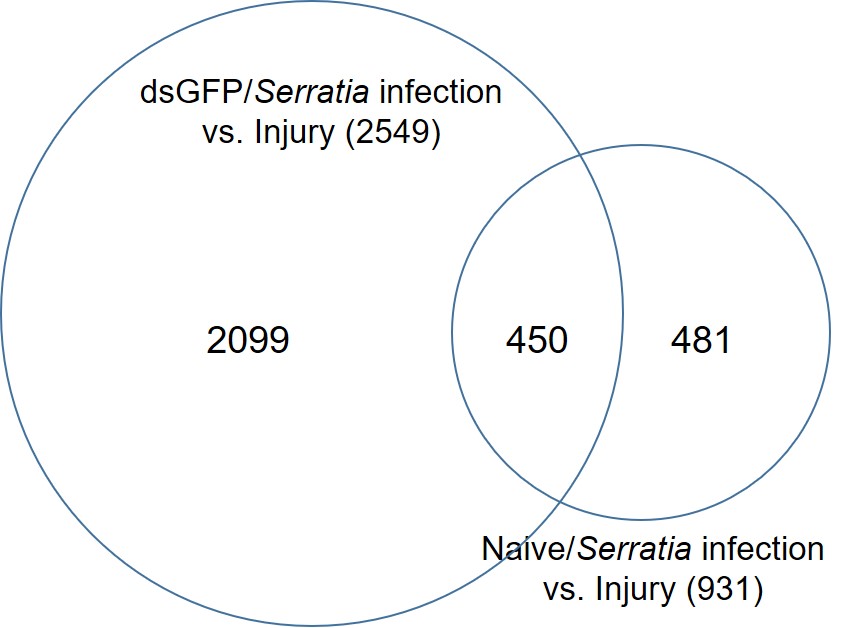

### Fig S5

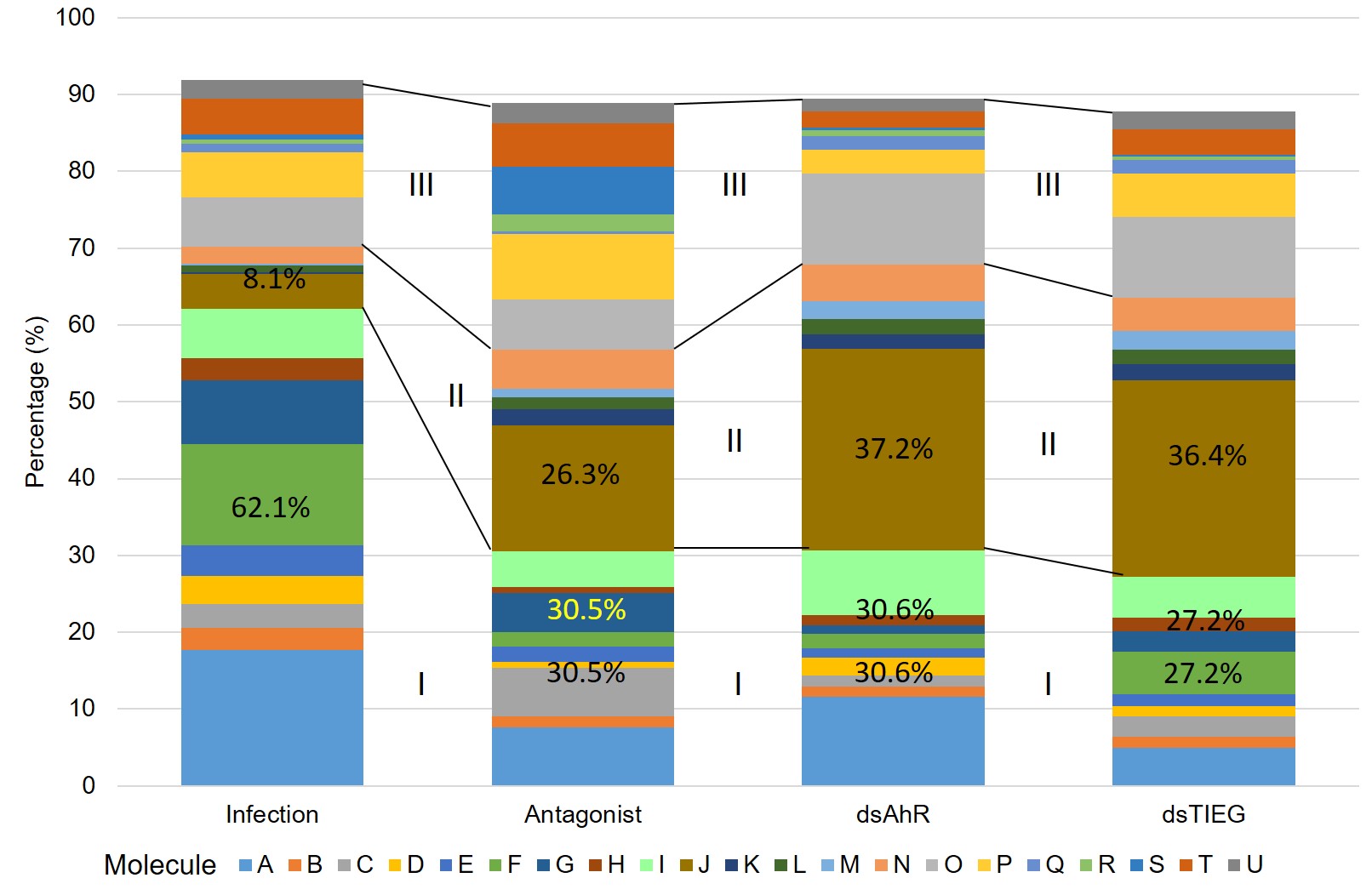
